# Automodification of N-terminal serine residues in PARP2 impacts PARP2 release from DNA damage sites

**DOI:** 10.1101/2025.07.01.662498

**Authors:** Saurabh Singh Dhakar, Albert Galera-Prat, Jin Cai, Julia Preisser, Tiila-Riikka Kiema, Andreas Ladurner, Lari Lehtiö

**Affiliations:** Faculty of Biochemistry and Molecular Medicine, University of Oulu, Finland; Department of Physiological Chemistry, Biomedical Center (BMC), Faculty of Medicine, LMU Munich, 82152 Planegg - Martinsried, Germany; International Max Planck Research School (IMPRS) for Molecules of Life, 82152 Planegg - Martinsried, Germany; Biocenter Oulu, University of Oulu, Finland

## Abstract

ADP-ribosyltransferases PARP1 and PARP2 are involved in DNA repair mechanisms and play a major role in detecting DNA damage. PARP1/2 enzymes transfer the ADP-ribosyl moiety from NAD^+^ to DNA damage proteins, histones and to itself, activating the DNA repair cascade. Recent studies have shown that Histone PARylation Factor (HPF1) forms a joint active site with the catalytic domain of PARP1/2 and alters their target modification site from glutamate/aspartate towards serine. In this work, we have identified key serine residues within the N-terminus of full-length PARP2 that are the main targets of PARP1/2 automodification. We demonstrated this using site-directed mutagenesis, gel-based PARylation assays of automodification reactions, and by measuring the release of PARP2 from the DNA damage site using a fluorescence polarization assay. We show that in the presence of HPF1, PARP2 serine 8 and serine 73 are predominantly ADP-ribosylated and serine 8, which is present in both human PARP2 isoforms, is the major site for PARP2 automodification. Our results provide insight into the mechanistic role of the N-terminus of PARP2 in the PARylation-dependent release of PARP2 from DNA damage sites.

## Introduction

The ADP-ribosylation of proteins is a post-translational modification that regulates a variety of biological processes such as cell cycle regulation, transcription, and DNA damage repair [1]. There are 17 ADP-ribosyltransferases (ARTs) in humans with homology to Diphtheria toxin (ARTDs) [2]. These enzymes, PARPs and tankyrases, regulate a wide range of cell signaling and especially PARP1-3 are known to recognize DNA lesion sites and activate DNA repair cascades by transferring ADP-ribosyl moiety from NAD^+^ to PARP enzyme itself, and to histones [2–5]. The transfer of ADP-ribose moiety can happen in the form of a single unit, called mono-ADP-ribosyl (MAR), and can be extended to poly-ADP-ribose chains (PAR). PARP1 and PARP2 PARylate histone tails, leading to chromatin remodeling. They also automodify themselves as a regulatory mechanism for their DNA binding affinity and activity [6]. Aspartates and glutamates are the automodification sites in PARPs, but in the presence of Histone Parylation Factor (HPF1) the activity is shifted to serines [2,7–10]. HPF1 forms a joint active site with the catalytic domain of PARP1/2 to modulate their activity [11,12]. It has been proposed that PARP1/2 automodification increases the repulsion between DNA and negatively charged PAR chains leading to their release from DNA [13]. The PAR chain is hydrolyzed by two enzymes known as poly (ADP-ribose) glycohydrolase (PARG) and ADP-ribosylhydrolase 3 (ARH3). PARG consists of both endo and exo-hydrolase activity between two ADP-ribose but cannot hydrolyze the last ADP-ribose moiety attached to the target protein residue. This ADP-ribose moiety can be removed by hydrolases with specificity to the residue that the MAR is attached to. For serine residues the hydrolase ARH3 is the enzyme that removes MAR [14–16].

PARP2 is a multi-domain protein that contains an unstructured N-terminal region, a WGR domain, and a C-terminal catalytic fragment consisting of a regulatory helical domain and the ARTD domain (**Fig. 1A**). The WGR domain of PARP2 recognizes the DNA break, while the ART domain is responsible for the catalytic activity and the helical domain is an autoinhibitory structure [17,18]. It has been shown that the N-terminus helps PARP2 in binding with DNA [3,18–20]. PARP2 has two isoforms in humans. Isoform-1 (Uniprot id: Q9UGN5-1) is a 583 amino acid protein, while isoform-2 (Uniprot id: Q9UGN5-2) lacks 13 amino acids (68 aa to 80 aa) in the N-terminal region (**Fig. 1B**).

**Figure 1.**
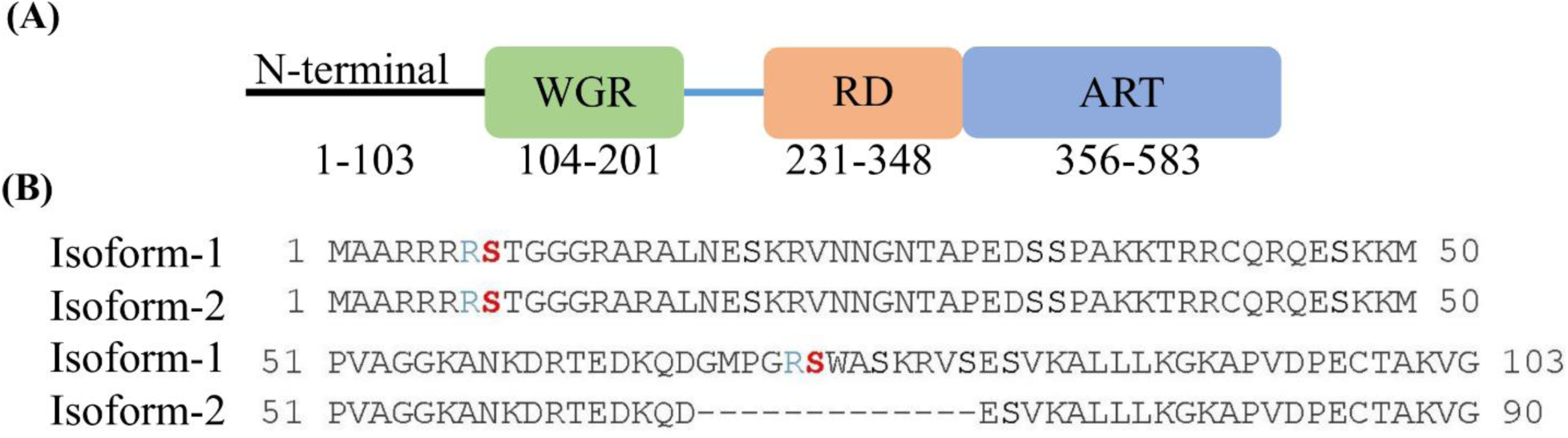
Full-length PARP2 domain arrangement and N-terminal sequence. (A) Structural domains of PARP2 and (B) Sequence comparison of the N-terminal disordered region between isoform 1 (accession: NP_5475) and isoform 2 (accession: NP_001036083) of PARP2. Based on mass-spectrometric analysis potential automodification serines 8 and 73 are highlighted in red while arginines upstream to serines are marked in blue.

Mass-spectrometry-based proteome studies in genotoxically stressed cells have shown that there are multiple targets for the post-translational PARylation of cellular proteins, but the majority of the ADP-ribosylation modification takes place at serine residues that follow a lysine or arginine residue [21–23]. In PARP1, three serine residues have been reported to be ADP-ribosylated. These predominantly account for the HPF1-dependent automodification of PARP1 during its enzymatic activity [24,25]. These serine residues are present in a disordered region (494 to 522) that precedes the WGR domain (541-637) and are also adjacent to lysine residues.

Based on these MS-based ADP-ribosylation sites and serine residues identified in PARP1, we investigated the presence of serine residues following a basic amino acid in the N-terminal of PARP2. In PARP2 isoform-1, there are two serines, Ser8 and Ser73, while in isoform-2 only Ser8 is present after the basic amino acid arginine (**Fig. 1B**). In MS investigation, Ser 8 and Ser73 in PARP2 have been detected to be ADP-ribosylated [21]. To understand the role of these serines in PARP2 automodification, we generated serine to alanine mutants in PARP2 isoform-1 that contains both putative modification sites and performed biochemical, biophysical and cell-based experiments. In this work, we show using site directed mutagenesis that N-terminal serines are the major automodification sites in PARP2. Using fluorescence polarization assays, we were able to test the effect of serine mutants on PARP2 release from DNA. We also carried out laser microirradiation assays in cultured cells to assess whether the mutated proteins were recruited or released from DNA damage sites in an altered manner, but could not readily determine an altered behaviour. This might be because other factors could impact recruitment and release kinetics in cells. The *in vitro* data identifies Ser8 as the main auto-ADP-ribosylated residue. In the longer canonical isoform (PARP2 isoform-1), Ser73 also contributes to automodification and PARP2 release from the DNA damage site.

## Results and Discussion

### Design and production of the serine mutants

To study the effect of serine residues in the automodification of PARP2, serine mutations were made to PARP2 isoform-1. Serines were mutated into alanine at position 8 (S8A, PARP2^S8A^), 73 (PARP2^S73A^), both 8 and 73 (PARP2^S8,73A^), and all 9 serines present at N-terminal region into alanine (PARP2^SA^). Two constructs were generated by inserting the serines at position 8 (PARP2^8S^) and 73 (PARP2^73S^) over no serine mutant (PARP2^SA^) to observe the recovery of the PARylation of PARP2 on the residues. The recombinant full-length PARP2 proteins were expressed in *E. coli* and purified with 4 step purification method. The serine mutations do not affect the protein expression, and 1.5-2 mg/L protein yield was obtained. These wild-type and serine mutant proteins were used to study the effect of the N-terminal serine residues on the auto-PARylation of PARP2.

### Analysis of the automodification of the serine mutants

To study the effect of serine residues on the automodification of PARP2, the activity assay for full-length wild-type PARP2 and serine mutants was performed using SDS-PAGE (**Fig. 2**). In the activity assay, different constructs of PARP2 were mixed with oligo (nicked double-strand dumbbell DNA containing 5’-phosphate), and HPF1 (as per reaction conditions), and the reactions were initiated by addition of NAD^+^. In the presence of oligo only, wild-type PARP2 did not show any apparent PARylation and was not affected by the presence or absence of HPF1 (**Fig. 2A, lanes 2, 7**). In the presence of NAD^+^ only, wild-type PARP2 showed a very low level of basal activity, but addition of oligo resulted in increased PARP2 activity, reflected in the form of a smear on gel (**Fig. 2A, lanes 3, 4**). The addition of HPF1 in the reaction containing oligo, PARP2 and NAD^+^ resulted in a drastic increase in wild-type PARP2 modification (**Fig. 2A, lane 5**). When both HPF1 and NAD^+^ were in the reaction, even in the absence of oligo, wild-type PARP2 automodified efficiently and produced more homogenous lower molecular weight PAR smear (**Fig. 2A, lane 6**). This can be because under *in-vitro* conditions, high protein concentrations would force the interaction of HPF1 and PARP2, which leads to the activation of PARP2 and results in its automodification. The high molecular weight smear from PARylated PARP2 was removed from acidic residues by hydroxylamine treatment but when HPF1 was present in the PARylation reaction, automodification was not eliminated (**Fig. S1**). This indicates that in the presence of HPF1, PARP2 modification occurs on serine residues.

**Figure 2.**
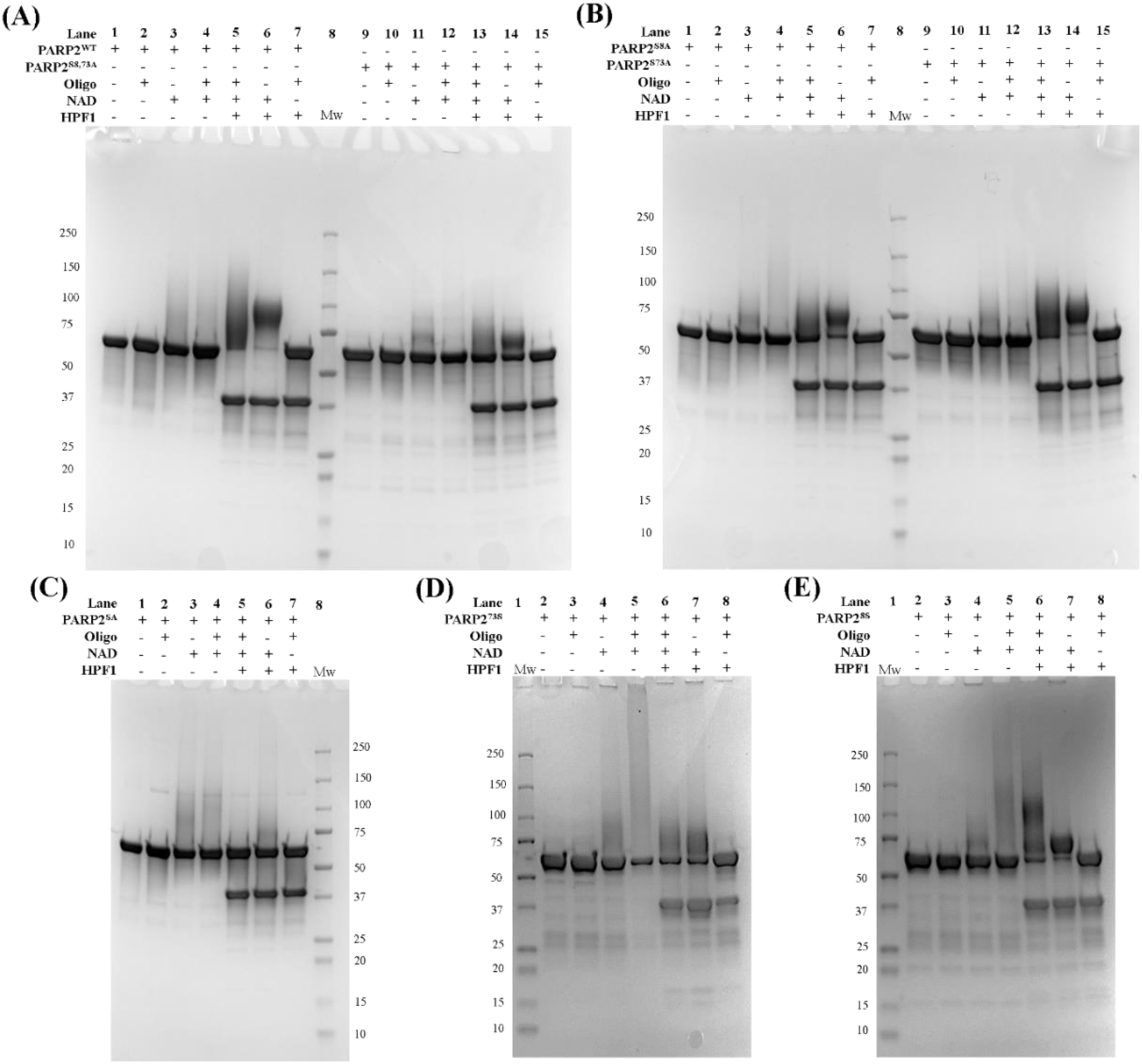
SDS-PAGE analysis of PARylation to visualize the effect of serine mutation on full-length PARP2 automodification: (A) wild-type PARP2 (lanes 1-7), PARP2^S8,73A^ mutant (lanes 9-15); (B) PARP2^S8A^ mutant (lanes 1-7), PARP2^S73A^ mutant (lanes 9-15); (C) PARP2^SA^ mutant; (D) PARP2^73S^ mutant; (E) PARP2^8S^ mutant.

In the absence of HPF1, all serine mutants showed a similar smear on gel as wild-type PARP2 (**Fig. 2A, lanes 2,3,4,10,11,12; Fig. 2B, lanes 2,3,4,10,11,12; Fig. 2C, lanes 2,3,4; Fig. 2D, lanes 2,3,4; Fig. 2E, lanes 2,3,4**). In the presence of HPF1, wild-type PARP2 showed a higher amount of smear as compared to serine mutants, while a fraction of serine mutant protein remained unmodified (**Fig. 2B**, **lane 5; Fig. 2B, lane 13; Fig. 2A, lane 13**). When all N-terminal serine residues were mutated into alanine (PARP2^SA^), most of the protein remained unmodified (**Fig. 2C, lane 5**). In the case of PARP2^S8A^, the amount of unmodified protein was higher than that of PARP2^S73A^, which indicates that Ser8 would be the main automodification site of PARP2 (**Fig. 2B**, **lanes 5,13**). PARP2^S8,73A^ mutant resulted in the loss of PARylation and most of the protein remains unmodified (**Fig. 2A, lane 13**). To verify the effect of serine mutations, Ser8 (PARP2^8S^), and Ser73 (PARP2^73S^) were inserted back into PARP2^SA^. In the presence of HPF1, the insertion of Ser73 (PARP2^73S^) increased PARylation as compared to PARP2^SA^ but most of the protein remained unmodified (**Fig. 2D, lane 6**), while the insertion of Ser8 (PARP2^8S^) resulted in the modification of majority of PARP2 (**Fig. 2E, lane 6**). The comparative analysis of the PARylation pattern among various serine mutants of PARP2 indicates that PARP2 Ser8 is the major automodification site, and Ser73 complements this function in the longer PARP2 isoform.

The effect of serine residues on automodification of PARP2 was corroborated with western blot analysis (**Fig. 3**). In the absence of HPF1, wild-type PARP2, and serine mutants showed similar PARylation patterns on the blot (**Fig. 3, lanes 2,4,6,8,10**). The addition of HPF1 changed the PARP2 automodification smear and the amount of unmodified protein. In the presence of HPF1, no amount of wild-type PARP2 remains unmodified (**Fig. 3, lane 3**). PARP2^S8A^ resulted in a significant amount of unmodified protein (**Fig. 3, lane 5**), while no unmodified protein was detected in PARP2^S73A^ (**Fig. 3, lane 7**). PARP2^S8,73A^ showed an increased amount of unmodified protein in comparison to PARP2^S8A^ (**Fig. 3, lane 9**), indicating Ser73 contributes to the automodification while majority of PARP2^SA^ remains unmodified (**Fig. 3, lane 11**). The insertion of Ser8 and Ser73 over PARP2^SA^ restored the automodification (**Fig. 3, lanes 14 & 16**). The western blot results also support that Ser8 of PARP2 is the major site for the automodification and Ser73 complements the automodification of PARP2 in its PARylation activity.

**Figure 3.**
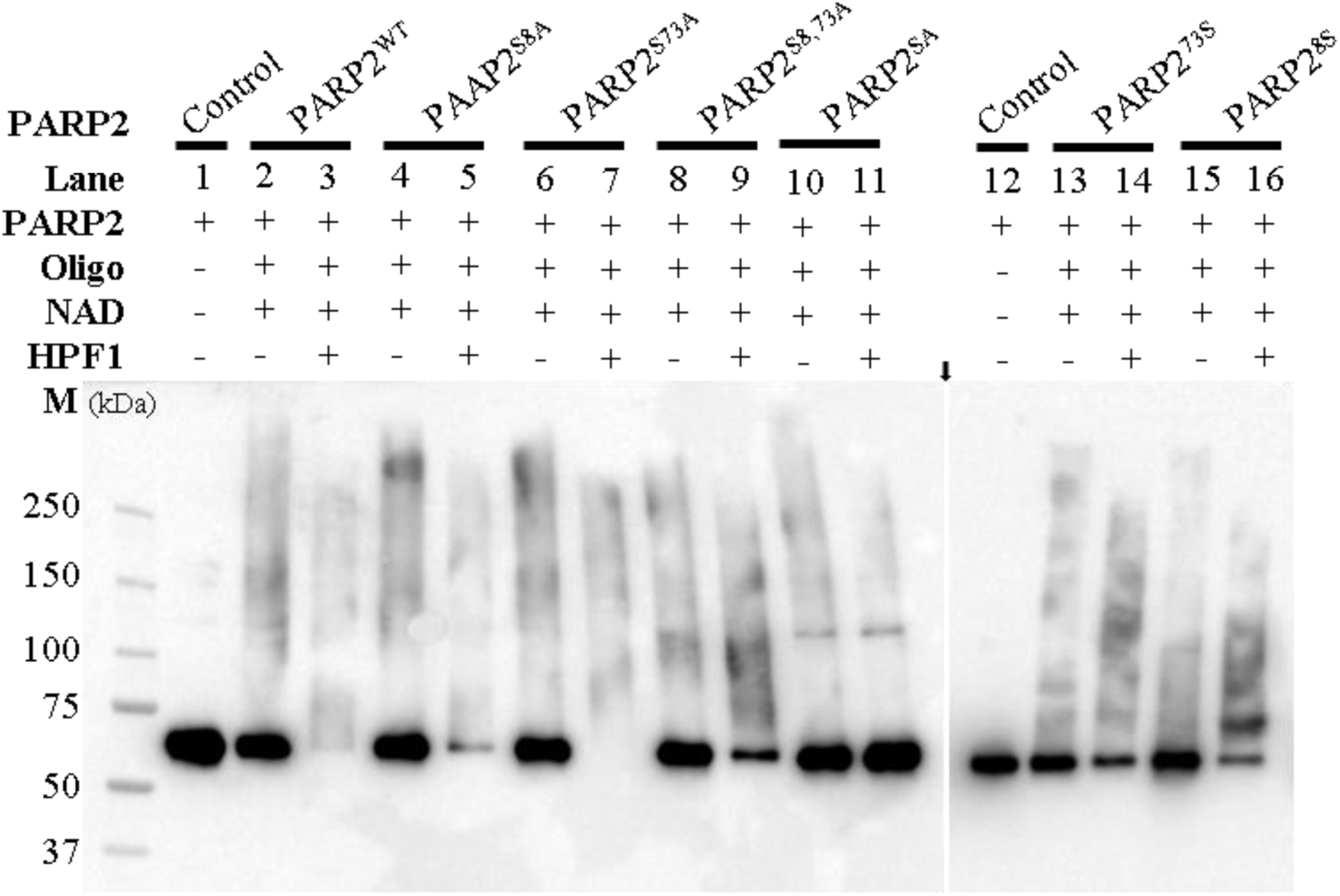
Western blot analysis of PARP2 activity in the PARylation assay to measure the effect of serine mutation on its automodification. The arrow between lanes 11 & 12 indicates the separation of two blotting membranes.

### PARP2 release from DNA upon automodification

In the activity assay, serine mutants showed a reduction in PARP2 automodification as compared to wild-type PARP2. We thus hypothesized that serine automodifcation by PAR could affect the release of PARP2 from the DNA it associates with. We therefore investigated the automodification-dependent release of PARP2 from DNA lesions using fluorescence polarization (FP) assays with labelled DNA oligonucleotide, thus mimicking DNA damage. The affinity of PARP2 to the oligonucleotide (fluorescein labeled nicked double-strand DNA with a 5’-phosphate) was 4.8 ± 2.9 nM (**Fig. S2**). For the automodification release experiments, we used a saturating concentration of 200 nM PARP2. The addition of 200 nM PARP2 to 5 nM oligo-DNA resulted in an increase in FP signal, indicating binding of PARP2 with the oligo (**Fig. S3**). After addition of 75 µM NAD^+^ in the reaction, FP signal started to decrease as PARP2 started to PARylate itself. Further, in the presence of HPF1, PARP2 automodification increased drastically (**Fig. 2**). To understand the effect of HPF1 on PARP2 release from DNA, 200 nM HPF1 was pre-mixed with 5 nM oligo-DNA and 200 nM PARP2, and the reaction was initialized again by adding 75 µM NAD^+^. In the presence of HPF1, wild-type PARP2 release from DNA was faster compared to the absence of HPF1 (**Fig. S3**). This agrees with the observation that HPF1 increases PARP2 automodification activity.

In our FP measurements, the addition of NAD^+^ in the reaction resulted in an initial increase in FP signal and only after a few minutes the signal started to decrease (**Fig. 4**). In the presence of HPF1, the increase in the FP signal was higher as compared to conditions without HPF1. We hypothesized that the increase in the initial FP was due to the auto-PARylation of PARP2, whereby the enzyme requires a certain amount of PAR to release from DNA, resulting in the initially higher polarization signal due to the increased molecular weight of the complex. After a certain amount of PAR formation, PARP2 molecules would then start releasing from the DNA. A similar effect of an increase in FP following the addition of NAD^+^ has also been reported for PARP1 [26]. To investigate the contribution of PAR to the initial increase in FP signal, we added the PARP2 catalytic inhibitor EB-47 [27] before initiating the PARP2 catalytic reaction with NAD^+^. The addition of EB-47 resulted in no changes in FP signal, i.e. neither an increase, nor a decrease (**Fig. S4**). This demonstrates that both the initial increase and the subsequent release from DNA depend on PARP2 activity.

**Figure 4.**
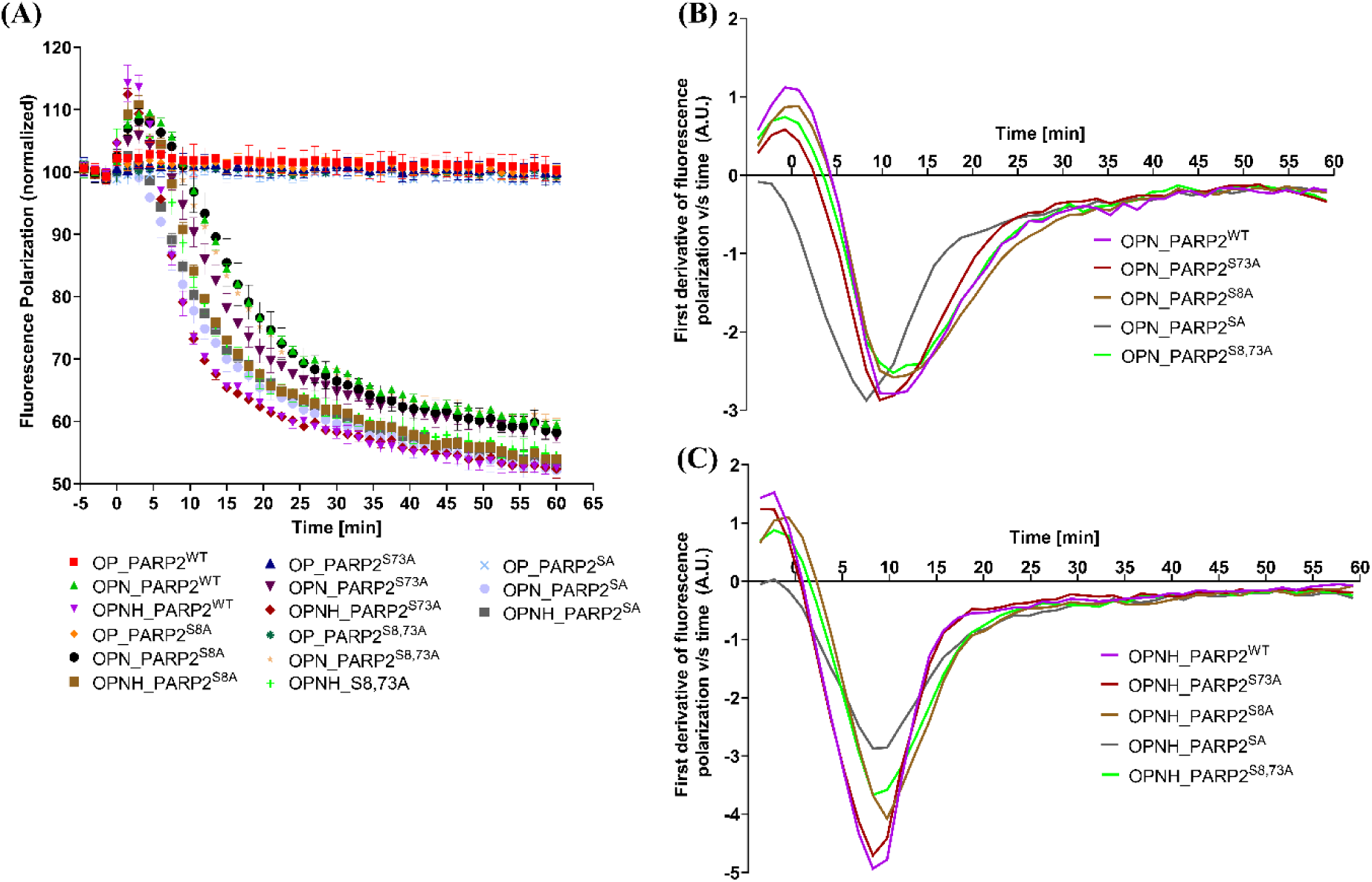
Representative PARP2 release from DNA monitored by fluorescence polarization (FP). (A) Time course FP measurement (every 90 seconds data points) to monitor PARP2 release from DNA. Reaction was initiated by adding NAD^+^ and at t=0 minute is the first FP measurement after adding NAD^+^. For comparative analysis among various PARP2 constructs, FP signal was normalized considering Oligo + Protein (OP) as 100 (maximum FP) and Oligo alone as zero (minimum FP). First derivatives of FP signal with time to compare the effect of serine mutations on release rates of PARP2: (B) without HPF1; (C) in presence of HPF1. [O: Oligo; OP: Oligo + PARP2; OPN: Oligo + PARP2 + NAD^+^; OPNH: Oligo + PARP2 + NAD^+^ + HPF1].

It should be noted that these experiments were carried out under saturating conditions, where there is an excess of unbound and unmodified PARP2. The unmodified PARP2 molecules could compete with the automodified PARP2 molecules to bind with DNA. As the reaction progresses, the amount of PARylated PARP2 molecules increases and results in a net decrease in the polarization signal.

### Effect of serine mutations on PARP2 release from DNA

Next, we assessed the DNA binding and dissociation kinetics using FP for PARP2 serine mutants in order to establish the effect of the mutations in the presence and absence of HPF1 (**Fig. 4A**). Because of the different amounts of the initial increase in FP signal and saturating protein concentration, it was not possible to directly compare the release among the PARP2 constructs. The difference was more evident when maximum release rates of PARP2 were calculated using the first derivatives of the FP signal vs time (**Fig. 4 B & C**). The lower (higher negative) value of the first derivative indicates the higher release rate, while the higher (less negative) value indicates the slower rate of release of PARP2 from DNA.

Based on comparative analysis of the first derivatives, we observed that in the absence of HPF1, all PARP2 constructs including wild-type and serine mutants released at similar rates (**Fig. 4B**). In contrast, when HPF1 was present, all PARP2 constructs except PARP2^SA^, released faster in comparison to conditions without HPF1. Since PARP2^SA^ does not contain any serines at N-terminus, our result highlights the role of the N-terminal serines in PARP2 automodification and in PARP2 release dynamics. In the presence of HPF1, PARP2^WT^ showed the highest release rate, followed by PARP2^S73A^. PARP2^S8A^ and PARP2^S8,73A^ were released slower than the wild-type and PARP2^S73A^. This reiterates our results from the gel-based activity assays, whereby both PARP2 Ser8 and Ser73 contributed to the automodification and subsequent release from DNA, with Ser8 as the key automodification site.

### Pre-steady state kinetics of automodification dependent DNA release

To precisely measure the effect of serine mutations on PARP2 release kinetics, we measured release rates through FP signals using stopped-flow kinetics in pre-steady state conditions. To perform the stopped-flow experiment, fluorescently labeled oligo (nicked double-strand DNA with a 5’-phosphate) was mixed with PARP2 and HPF1 in 1:1:1 ratio at a 200 nM concentration, achieving a saturated state, where all the PARP2 molecules are bound to DNA and the first-order rate kinetics can be monitored. To initiate the reaction, 500 µM NAD^+^ was added to the reaction using the stopped-flow syringe and FP signal was measured for all the PARP2 constructs (**Fig. 5A**). In this experiment, it was evident that the PARP2 variants released from the DNA with different rate (**Fig. 5B**). The PARP2^WT^ released fastest (0.09572 ± 0.00098 sec^-1^), followed by PARP2^S73A^ with a slower rate (0.08987 ± 0.00036 sec^-1^). PARP2^S8A^ had a significantly reduced release rate from DNA and the rate was almost half of that of the wild-type protein (0.05223 ± 0.00107 sec^-1^). PARP2^S8,73A^ showed the additive effect of the S73A mutation and resulted in an even slower rate of release (0.04606 ± 0.00118 sec^-1^). PARP2^SA^ released with slowest rate (0.02742 ± 0.00146 sec^-1^) among all the PARP2 constructs, indicating that additional serine residues at the N-terminus of PARP2 contribute to the release kinetics (**Fig. 2**). Overall, we found that there appears to be a direct correlation with the amount of PARP2 automodification on N-terminal serine residues and the release rate from DNA.

**Figure 5.**
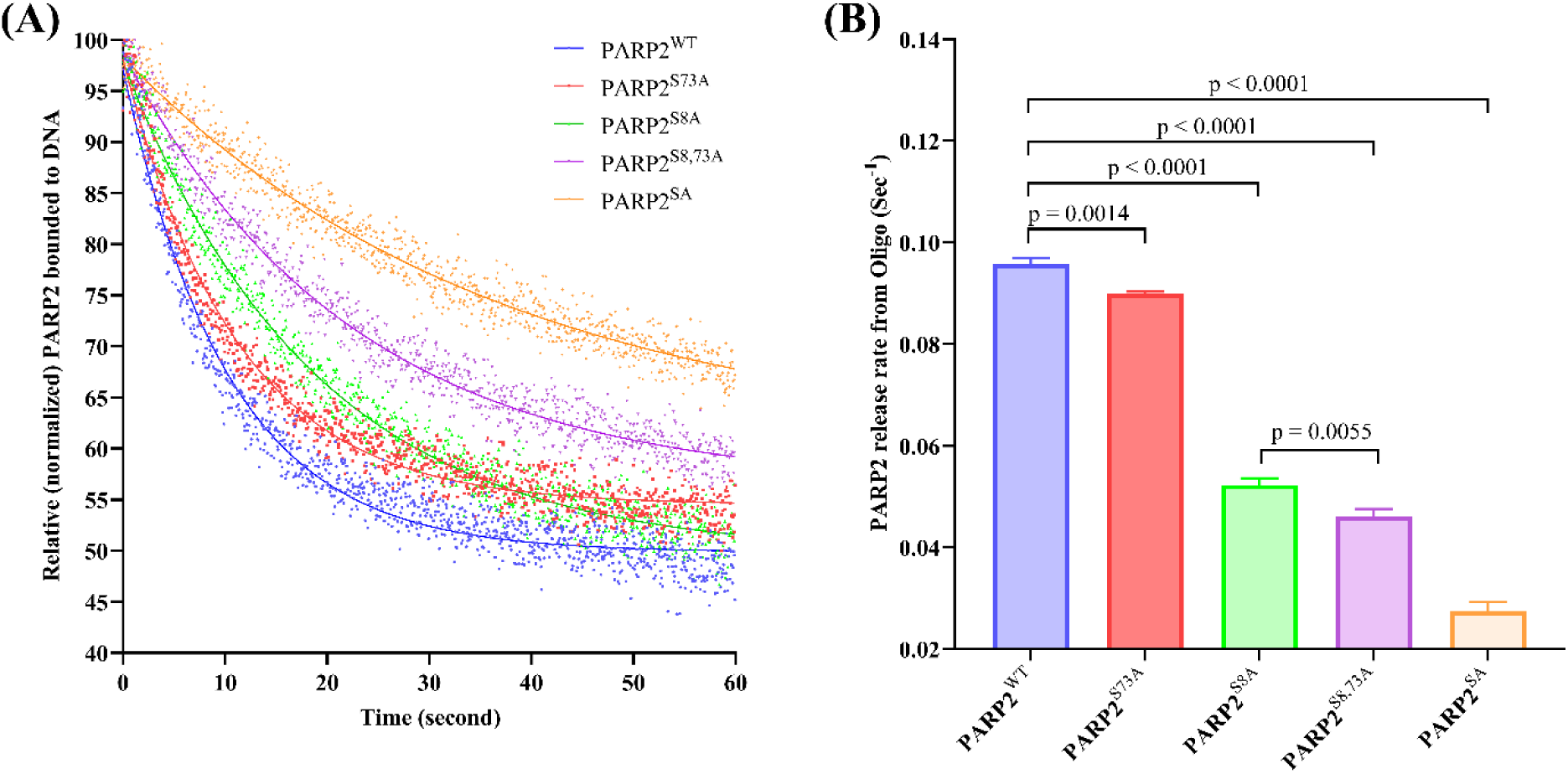
Time-course fluorescence polarization measurement of PARP2 wild-type and serine mutants using stopped-flow measurements. (a) PARylation-dependent PARP2 release from DNA. (b) Statistical analysis of release rate kinetics for PARP2 serine mutants. PARP2 release rates (sec^-1^) shown are average ± s.d. from three independent experiments with internal replicates.

### DNA release dynamics of PARP2 in live cells

We next wanted to study the retention and release rates of PARP2 and test the effects of the serine mutations in the DNA damage context using live-cell imaging with laser microirradiation, specifically by following the dynamics of fluorescently labelled PARP2. PARP1 is the first line of defense upon DNA damage, which also affects the mode of PARP2 recruitment at DNA lesions [28,29]. To exclude the interference from endogenous PARP1/2 in the recruitment of the fluorescently-tagged PARP2 enzyme, we used PARP1/2 double knockout U2OS cell lines in the experiments. PARP2 constructs including wild-type and all PARP2 serine mutants (PARP2^S8A^, PARP2^S73A^, PARP2^S8,73A^, and PARP2^SA^) were transfected in PARP1/2 double-knockout U2OS cell lines. Upon induction of DNA damage using the laser, all PARP2 constructs recruited to the induced DNA lesion site. During time-course imaging, the intensity of recruited PARP2 at the lesion site started to gradually decrease over time (**Fig. 6A**). Based on the analysis of fluorescence intensities at the DNA damage site, we could not observe a difference in the retention time for all tested PARP2 constructs (**Fig. S5**). Since the U2OS cells also express PARG, which efficiently hydrolyzes PAR in the cell and therefore could affect the PAR dependent release of PARP2 from DNA lesion site, we also conducted experiments with PARG inhibitor to exclude this possibility. We treated cells with 1 µM PARG inhibitor PDD00017273 (Sigma) during the microirradiation experiment and imaging. However, we did not observe any differences in the release kinetics of our tested PARP2 serine mutants from DNA damage sites (**Fig. 6B**). This indicates that PARP2 release from DNA lesion sites is not solely dependent on the automodification of the N-terminal serines of PARP2.

**Figure 6.**
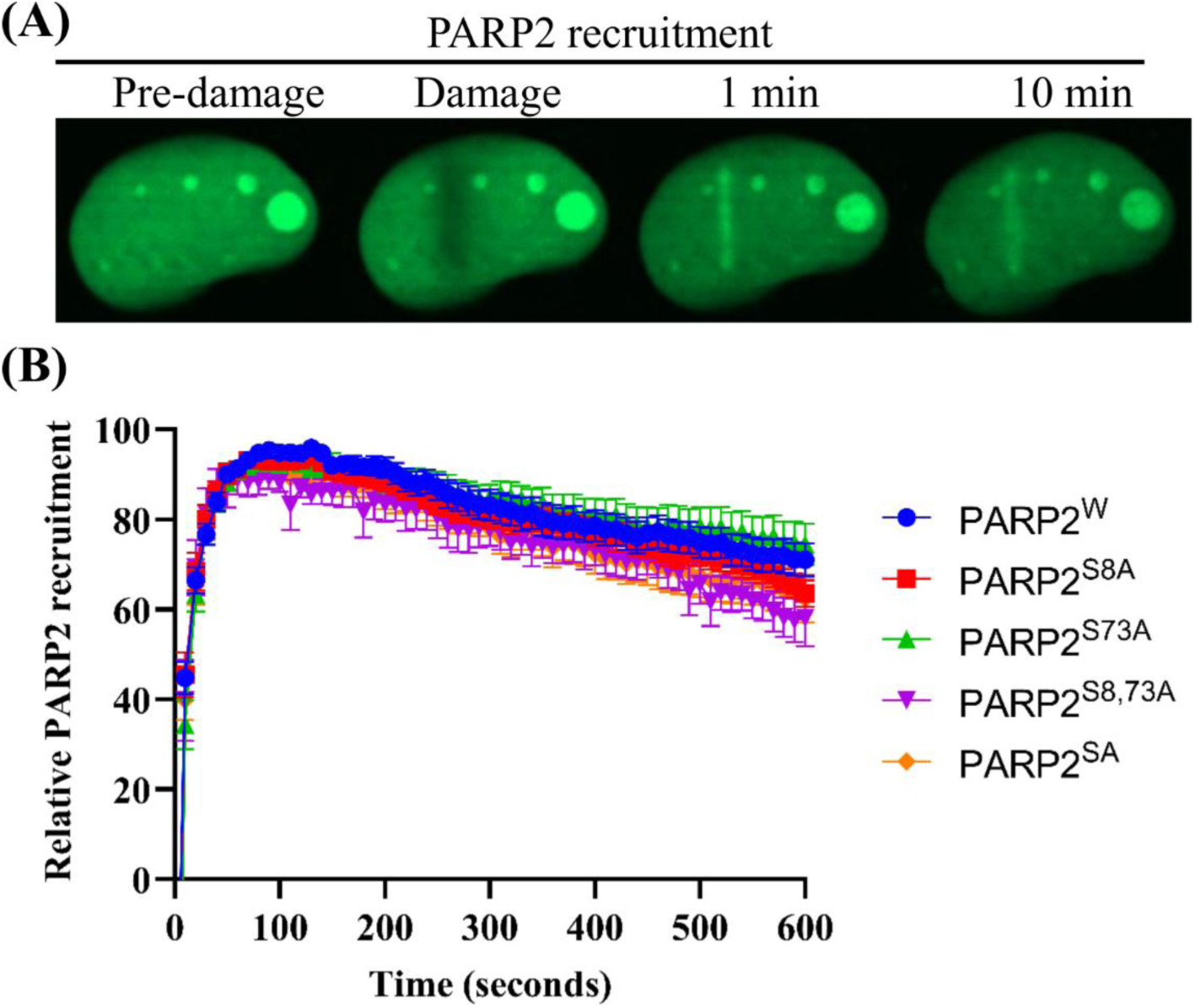
PARP2 recruitment and release from DNA lesion sites in PARP1/2 double knockout U2OS cells. (A) Representative images of GFP-PARP2 enrichment after DNA damage during time course imaging. (B) Relative kinetic recruitment and release of GFP-PARP2 from DNA lesion in the presence of 1µM PARG inhibitor. Data shown as mean ± SEM (n > 10 per construct) at the microirradiation site. Release for PARP2 wild-type and serine mutants were compared by normalizing the GFP intensities for every cell considering pre-damage intensity as zero and maximum GFP intensity to 100.

Apart from automodification of PARP2, other factors such as histone modification, chromatin remodeling, recruitment of DNA repair proteins might be involved in facilitating PARP2 release from DNA lesions. In our PARylation gel assay and the FP experiment, we observed that mutation of Ser8 and Ser73 in PARP2 resulted in a reduction of PARP2 automodification, which in turn lowered its release from DNA. These serine mutations do not completely abolish PARP2 activity, nor do they promote the retention or trapping of PARP2 to DNA (at least not as strongly as PARP inhibitors do). This is in line with the study showing that HPF1 would not be obligatory for DNA damage repair to occur and that modification on glutamate and aspartate may compensate for HPF1 loss [30]. As Mahadevan *et al.* described, there are freely diffusing and DNA bound PARP molecules in the DNA damage region [31] and this heterogenous population of PARP2 could limit our ability to detect differences between wild-type and serine mutant PARP2 release in the cellular conditions due to the limited imaging resolution.

### Structural basis of N-terminal serine preference for automodification

As we had identified, Ser8 emerged as the preferential PARylation auto-modification site of PARP2, so we attempted to generate a model of the ternary complex of PARP2 ART domain, HPF1, N-terminal peptide and substrate NAD^+^. Alphafold 3 [32] generated a model where the PARP2 and HPF1 complex are in good agreement with the experimentally determined structure (PDB id: 6TX3) [12] (**Fig. S6 A,B**). NAD^+^ bound the ART domain similarly to the experimental structure of PARP1 with an unhydrolyzable analog benzamide adenine dinucleotide (BAD) (PDB id: 6BHV) [33] (**Fig. S6 C,D)**. The 15-mer peptide (aa 1-15, MAARRRRSTGGGRAR) from the N-terminus with Ser8 at the center is positioned in the model in such a way that the positively charged Arg7 is placed in a negatively charged patch of HPF1 (**Fig. 7A**) and the serine side chain is positioned within a 3.6 Å distance from the O3’ of NAD^+^ ribose sugar (**Fig. 7B**). For the Ser8 peptide, there are four consecutive arginine residues before the serine, and these could contribute to the efficient recruitment of the peptide to the peptide binding cleft formed by the PARP2-HPF1 complex. Other peptides of similar lengths (**Table S1**) were modelled with PARP2-HPF1-NAD^+^ complex and found that with a low confidence in modelling Ser73 is also located near the NAD^+^ and active site (**Fig. S7**). Other N-terminal peptides with serine residues did not produce any similar models. Based on this serine 8 is structurally preferred for automodification during the catalytic mechanism of PARP2.

**Figure 7.**
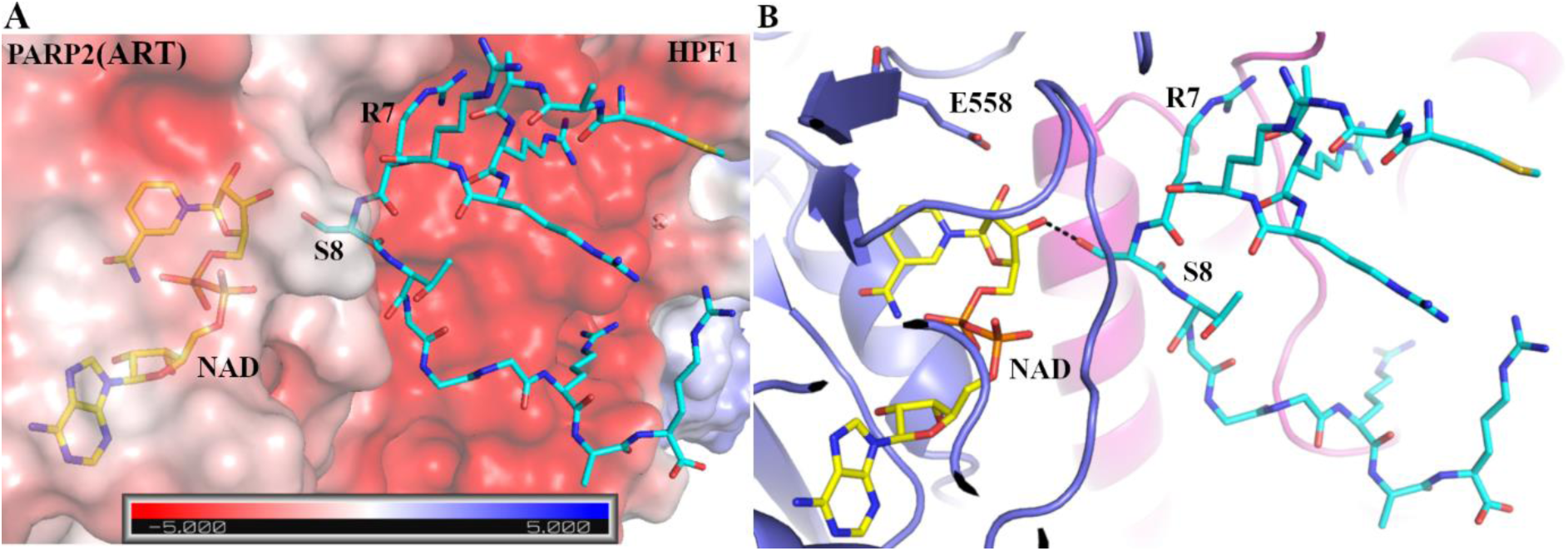
Alphafold 3 model of PARP2-HPF1-NAD^+^ with N-terminal peptide complex. (A) Arginine upstream to Ser8 in peptide (cyan) interacts with the negatively charged (red) patch in HPF1 [shown as gradient of red (negative charge) to blue (positive charge)]. Surface electrostatic charge is calculated using APBS-PDB2PQR software [34]. (B) 15-mer peptide (cyan) interaction at PARP2 ART (blue)-HPF1 (magenta) where Ser8 is oriented near ribose sugar of NAD^+^ (yellow) within 3.5 Å distance.

### Conclusions

The N-terminal region is an unstructured part of the PARP2 enzyme. Beyond its high affinity for DNA [20,35], its functional role is not understood. In the context of DNA damage, serines in PARP2 are the main ADP-ribosylated residues in the presence of HPF1 [7]. Here we focused on these identified PARP2 serine residues in the N-terminal region of the enzyme by mutating them to understand the impact of these residues on automodification and the subsequent release from DNA. Together, our PAR activity assays and FP based binding assays showed that mutations of Ser8 or Ser73 to alanine decreased PARP2 automodification and caused a delay in release of PARP2 from DNA. Ser8 affected the automodification and reduced the release rate of PARP2 more than Ser73, suggesting that Ser8 plays a major role in automodification during PARP2 catalytic mechanism. This is in agreement with activity assays showing Ser8 to be the main ADP-ribosylation site in the biochemical assay. Ser8 is present in both human PARP2 isoforms and the role of Ser73 in the longer isoform (PARP2 isoform-1) appears to be enhancing or complementing the function of Ser8. However, we did not observe differences in the release of the PARP2 constructs from DNA damage created using microirradiation in live cells. This suggests that PARP2 release from the region of DNA damage may not solely depend on HPF1-mediated PARP2 automodification, but other factors such as histone modifications, chromatin remodeling and the recruitment of DNA repair proteins might be involved in facilitating the release of PARP2. The PARP2 mutant without any N-terminal serines also released from DNA in the biochemical assay and it is possible that the effect of serine mutations on PARP2 release is not large enough, making the quantification in this established cellular assay challenging. In sum, our biochemical and biophysical data revealed a key role of Ser8 in PARP2 automodification and shed light on the overall mechanistic role of the disordered N-terminal region of PARP2 during DNA damage repair.

## Materials and methods

### Cloning of serine mutant in full-length PARP2 and full-length HPF1

The constructs for expression of codon-optimized full-length PARP2 isoform-1 (residues 1-583, UniProt ID: Q9UGN5) and full-length HPF1 (residue: 1-346, Uniprot ID: Q9NWY4) were previously described [18,35]. Inserts with serine mutations in the wild-type full-length PARP2 were generated using site-directed mutagenesis and by assembly PCR. To make an expression construct for full-length PARP2 Serine mutants, inserts were cloned into pNH-Trxt plasmid (addgene # 26106) using SLIC cloning method [36]. The pNH-Trxt plasmid was linearized in PCR using site-specific primers. 100 ng linearized plasmid was mixed with gene PCR products in a 1:3 molar ratio and incubated with T4 DNA polymerase for 2–3 min at room temperature. The mixture of insert and plasmid was transformed into *E. coli* strain NEB5α cells. The colonies were grown at 37°C overnight on LB agar media plates containing 10 mM benzamide as PARP2 inhibitor to overcome PARP2 toxicity for *E. coli*, 50 µg/ml kanamycin as an antibiotic, and 5% sucrose for SacB-based negative selection [35,37,38]. PARP2 constructs contain “His6 tag-TEV protease site-full length PARP2” and HPF1 contains His6 tag-TEV protease site followed by HPF1 coding sequence. For a mammalian cell-based study, PARP2 inserts including wild-type and serine mutants were cloned into EGFP-C1 vector, where constructs contain GFP at N-terminal to full-length PARP2.

### Protein expression of HPF1, wild-type and serine mutants of PARP2

Plasmid containing full-length PARP2 (wild-type or serine mutants) and full-length HPF1 were transformed into *E. coli* strain BL21(DE3) cells for protein expression and incubated at 37 °C overnight. Transformed colonies from overnight grown plates were inoculated in 500 ml autoinduction media and 0.8% (w/v) glycerol, 50 μg/ml kanamycin antibiotic were added to it. 10 mM benzamide was supplied to PARP2 culture medium [37]. *E. coli* cultures were incubated at 37 °C until an OD600 of 1.2, and then incubated for 18 h at 16 °C. Next day, cells were harvested by centrifugation at 4200 × g for 45 min at 4 °C. Pellets were resuspended in lysis buffer (50 mM HEPES, 500 mM NaCl, 10 mM imidazole, 0.5 mM TCEP, 10% (w/v) glycerol, pH 7.5) and 100 µM protease inhibitor pefabloc [4-(2-Aminoethyl) benzenesulfonyl fluoride hydrochloride] (Sigma, ref: 11429876001) was added in it. Cells were resuspended and flash frozen in liquid nitrogen and stored at − 20 °C.

### Protein purification of HPF1, wild-type and Serine mutants of PARP2

All the constructs of full-length PARP2 and HPF1 were purified as described previously [35]. The HPF1 construct was purified using multi step purification *i.e.,* immobilized metal affinity chromatography (IMAC), Histidine tag cleavage by TEV protease, reverse IMAC, and size exclusion chromatography (SEC) while PARP2 purification consists of one additional step of ion exchange chromatography between IMAC and histidine tag cleavage. Harvested cells were thawed and DNase (2 µg/ml) was added to HPF1 but not in the PARP2 cell pellet. Cells were lysed by sonication and lysate was centrifuged at 39,000 × g, at 4 °C for 1 hr. Supernatant for HPF1 protein was filtered and loaded on lysis buffer equilibrated Ni–NTA column while PARP2 supernatant was incubated with Ni-NTA resins for 1hr at 4 °C. The column was washed with 3 column volumes of wash buffer 1 (30 mM HEPES, 1 M NaCl, 10% glycerol, 0.5 mM TCEP, 10 mM imidazole, pH 7.5) and wash buffer 2 (30 mM HEPES, 500 mM NaCl, 10% glycerol, 0.5 mM TCEP, 50 mM imidazole, pH 7.5). Protein was eluted in an elution buffer (30 mM HEPES, 500 mM NaCl, 10% glycerol, 0.5 mM TCEP, 250 mM Imidazole, pH 7.5).

The IMAC eluted fraction for PARP2 was loaded on 5 ml Heparin column which was pre-equilibrated in Heparin buffer A (30 mM HEPES, 300 mM NaCl, 10% glycerol, 0.5 mM TCEP, pH 7.5) and after binding of protein column was washed with the same buffer. Protein was eluted with a linear gradient of 50 ml heparin buffer B (30 mM HEPES, 1.5 M NaCl, 10% glycerol, 0.5 mM TCEP, pH 7.5). PARP2 protein containing fractions from heparin and the IMAC elution for HPF1 were digested with TEV protease in 10 kDa dialysis bag in dialysis buffer (30 mM HEPES, 250 mM NaCl, 10% glycerol, 0.5 mM TCEP, pH 7.5), for overnight at 4 °C. Overnight TEV digested sample was supplemented with 25 mM imidazole before loading to wash buffer 2 pre-equilibrated IMAC column. The column was washed with wash buffer 2 and the flow through was collected. The collected protein sample was loaded on pre-equilibrated (30 mM HEPES, 500 mM NaCl, 10% glycerol, 0.5 mM TCEP, pH 7.5) 16/600 superdex-200 (Cytiva) (for PARP2) or 16/600 superdex-75 (Cytiva) (for HPF1) size-exclusion chromatography column. Purified protein was concentrated using 30 kDa (for PARP2 protein) or 10 kDa (for HPF1 protein) membrane filter (Millipore). Concentrated protein was aliquoted in small volumes, flash-frozen in liquid nitrogen and stored at −70 °C. All steps of the protein purification were performed at 4 °C in a cold room.

### Activity assay of PARP2 wild-type and serine mutants on SDS-PAGE

The PARylation activities of PARP2 wild-type and serine mutants were performed on SDS-PAGE gel (BioRad) as described previously [18] using the reaction buffer containing 50 mM Tris, 5 mM MgCl_2_, pH 8.0 (prepared as 10 × concentration). To monitor the effect of serine mutation on the auto-modification of PARP2, 7.6 µM PARP2 (wild-type or serine mutant) was mixed with 7.6 µM nicked double stranded dumbbell DNA (5’p GGG TCT TTT GAC CCT CGA GCT TTT GCT CGA 3’) and incubated for 10 min at room temperature. 7.6 µM HPF1 was added to the DNA protein mixture and the PARylation reaction was initiated by adding 1 mM NAD^+^ to the protein mixture and reactions were incubated for 1 h at 25 °C. PARP2 activity was stopped by adding 1 mM Olaparib to the reaction. SDS loading buffer was added to the reactions. Samples were separated on the 4–20% SDS-PAGE gel at 120V for 1h and stained with page blue staining solution.

### Western blot assays

The activities for PARP2 constructs were performed using 1 µM oligo, 1 µM PARP2, 1 µM HPF1, and 500 µM NAD^+^. The reaction was incubated for 5 min at 25 °C and stopped using 500 µM Olaparib. SDS loading buffer was added to the reactions and samples were separated on the 4–20% SDS-PAGE gel at 120V for 1h. Protein samples were transferred on transblot turbo 0.2 µm Nitrocellulose membrane (BioRad) using semi dry transfer with BioRad TransBlot system at constant 10 W for 30 min. The blotted membrane was blocked in 5% skim milk in TBST (TBS with 0.1% tween-20) for 1 h at 25°C followed by washing with TBST for 2x 15 min. The membrane was incubated overnight at 4°C temperature with anti-PARP2 primary antibodies against internal amino acids of PARP2 (aa 402-415, LDLFEVEKDGEKE) (Invitrogen, ref: PA5-18468), 1:1000 diluted in TBST- 3% BSA. Next day, membrane was washed 2x 15 min with TBST and incubated with peroxidase conjugated Rabbit-Anti-Goat secondary antibodies (Jackson immunoresearch, article no.: 305-035-003), 1:5000 diluted in TBST-3% BSA for 1 hour. The blotted membrane was washed 2x 15 min times with TBST and imaged using peroxide solution (BioRad ECL substrate) in BioRad ChemiDoc XRS^+^ or Azure 200 western blot imager.

### PARP2 modification and release from DNA using fluorescence polarization

The fluorescence polarization (FP) assay was measured using a Tecan Infinity M1000pro plate reader. The measurement was done in 50 µl volume in 96-wells plates. The instrument was calibrated with 1 nM fluorescein solution in 10 mM NaOH using the theoretical polarization value of 27 mP for free fluorescein. The fluorescein was excited at 470 nm (5 nm bandwidth) and the emission was recorded at 520 nm (20 nm bandwidth) wavelength. Other parameters, including the number of flashes and signal settling time, were manually set to 50 and 100, respectively. To measure automodification dependent PARP2 release from DNA, 5 nM fluorescein labeled nicked double-strand DNA with a 5’-phosphate oligo (5’p GGA AGT CTT iFluor TTG ACT TCC TCG AAG CTT TTG CTT CGA 3’), 200 nM PARP2, 200 nM HPF1 were used in optimized fluorescence polarization buffer (10 mM HEPES, 150 mM NaCl, 0.1 mg/ml BSA, 0.1 mM TCEP, 0.1 mM EDTA, pH 8.0). The PARylation reaction initiated by adding 75 µM NAD^+^ (final concentration in the reaction) and polarization was measured. The FP measurements were carried out in the kinetic scan mode at 25 °C. The time interval between cycle and number of cycles were determined independently for the experiment, depending on the number of scanned wells. Polarization data was analyzed, and graphs were plotted using GraphPad Prism10.

### Pre-steady state kinetics of PARP2 release using stopped flow measurements

To measure the pre-steady state kinetics of PARP2 release from DNA, fluorescence polarization of the PARylation reaction was monitored using Kinet Asyst Stopped-Flow Spectrometer (TgK Scientific, SF-61DX2). The fluorescence polarization was measured at an excitation wavelength of 492 nm and emission using 520 nm (10 nm FWHM) parallel and perpendicular bandpass filters. To measure the polarization, 200 nM fluorescein labeled oligo (5’p GGA AGT CTT iFluor TTG ACT TCC TCG AAG CTT TTG CTT CGA 3’), 200 nM PARP2, 200 nM HPF1 and 500 µM NAD^+^ concentrations were used in fluorescence polarization buffer (10 mM HEPES, 150 mM NaCl, 0.1 mg/ml BSA, 0.1 mM TCEP, 0.1 mM EDTA, pH 8.0). To set up the reaction, 400 nM (2x concentration) of Oligo, PARP2, HPF1 were premixed in one syringe and the other syringe was filled with 1 mM NAD^+^ (2x conc). The PARylation reaction was initiated by mixing the contents of the two syringes. For each stopped-flow reaction, 4 shots for 60 seconds with 1024 data points were collected. PARP2 release rate from DNA was calculated using a formula (Y = - A * exp^(-R^ * ^X)^ + C) in Kinetic Studio 5. For every PARP2 construct, 3 independent polarization measurements (4 shots per measurement) were performed, and average rates of release were calculated. Normalized data for wild-type PARP2 and serine mutants plotted using GraphPad 10.

### Cell cultures and transfection

Human osteosarcoma U2OS cell line with PARP1/2 double knockout [PARP1/2 KO] were used for the expression of GFP labeled PARP2 constructs. Cells were grown at 37°C in the presence of 5% CO2 in Dulbecco’s Modified Eagle Medium (DMEM)(Gibco, ref: 31885049) supplied with final concentration of 10% Fetal bovine serum (FBS) (Gibco, lot: 10270106), 100 units / ml penicillin and 100 µg / ml streptomycin antibiotics (Gibco, ref: 15140-122). Around 20,000 Cells were seeded in 8-well Lab-Tek chambers (Thermo fisher scientific) and overnight grown cells were transiently transfected with GFP-PARP2 construct using XtremeGENE HP DNA transfection reagent (Sigma, ref: 06366546001). The transfected cells were incubated in DMEM media for protein expression and after 24 hrs used for microirradiation and live cell imaging.

### Live-cell imaging and DNA microirradiation

The imaging experiments were performed on a Zeiss confocal spinning-disk microscope. Transfected cells were maintained in Leibovitz L-15 media (GIBCO, ref: 21083027). In desired conditions, Leibovitz media was supplied with 1 µM PARG inhibitor (Sigma, ref: SML1781) and incubated for 1 h before imaging. Cells with a comparable level of GFP-PARP2 expression were selected for imaging. DNA damage was induced using 10% laser power (355 nm) for 1 sec exposure time using a single-point scanning head (Rapp Opto Electronics) and time-course dynamics of PARP2 enrichment at damage sites were monitored by live-cell imaging for 10 min in every 10 second.

### Image analysis of live-cell experiments

The GFP-PARP2 enrichment at microirradiation site was quantified using a custom macro as described by Blessing et al., 2020 [39] in Fiji/ImageJ using StackReg plugin [40–42]. The damaged region of interest (ROI) with 25x100 pixels was selected for fluorescence enrichment at the damage site and the cell nucleus was selected as the region for GFP-PARP2 expression in the nucleus. The background was defined as a region outside the nucleus. The PARP2 recruitment was analyzed by comparing the fluorescence intensity at the damage site before and after the microirradiation. PARP2 enrichment and release from the damaged site were calculated using the fluorescence intensity of ROI as “[(Damage -background) / avg (damage site before irradiation)] / [(Nucleus - background) / avg (nucleus before irradiation)]”. Using the measured fluorescence intensities from Fiji/ ImageJ, the graphs were plotted in GraphPad Prism 10 and PARP2 release from DNA damage site was analyzed.

## Author contributions

A.G-P., L.L. conceptualization; S.S.D., J.P., T-R.K. investigation; S.S.D., A.G-P, J.C. Formal Analysis; S.S.D. writing-original draft; S.S.D, A.G-P., J.C., J.P., T-R.K., A.L., L.L., writing-reviewing and editing; S.S.D, A.G-P., A.L., L.L., funding acquisition; A.G-P., A.L., L.L. supervision.

## Acknowledgements

This work was funded by Research Council of Finland (347026 for LL), (354387 for AGP) and the Deutsche Forschungsgemeinschaft (German Research Foundation; DFG; Project-IDs 213249687 – SFB1064 and 325871075 – SFB1309 for AL). The Boehringer Ingelheim Fonds and The Magnus Ehrnrooth Foundation provided travel grant support for research visits to LMU (for SSD). Biocenter Oulu Structural Biology core facility, member of Biocenter Finland, Instruct-ERIC Centre Finland and FINStruct, as well as of “Proteomics and Protein Analysis” and Sequencing core facilities are gratefully acknowledged. We thank Prof. Taina Pihlajaniemi for providing peroxidase conjugated Rabbit-Anti-Goat secondary antibodies.

## Notes

### Competing Interest Statement

The authors have declared no competing interest.

