## Supplementary information for "Automodification of N-terminal serine residues in PARP2 impacts PARP2 release from DNA damage sites"

CONTENT

**Figure S1.** Effect of hydroxylamine on automodification of wild type and serine mutants of full length PARP2.

**Figure S2.** Fluorescence polarization-based Kd determination of PARP2 with oligo.

**Figure S3.** Representative fluorescence polarization assay to monitor the effect of HPF1 on PARP2 release from DNA.

**Figure S4.** Fluorescence polarization assay to monitor the effect of PARP2 catalytic inhibitor EB-47 on its release from DNA.

**Figure S5.** Relative release of PARP2 from DNA lesion in PARP1/2 double knockout U2OS cells.

**Figure S6.** Structural analysis of AlphaFold 3 generated PARP2-HPF1 complex and $NAD^+$ binding.

**Figure S7.** Alpha-fold 3 model of PARP2-HPF1-$NAD^+$ complex showing Ser73 is oriented near PARP2 catalytic and $NAD^+$ binding site.

**Table S1.** Peptide sequences used for model generation in Alphafold3.

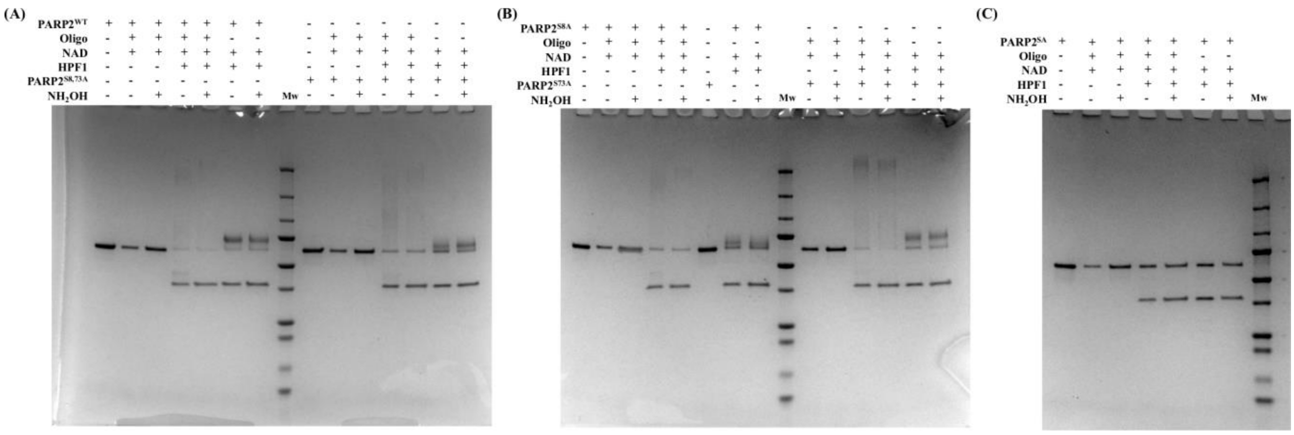


**Figure S1.** PARylation assay to visualize the effect of hydroxylamine on automodification of wild type and serine mutants of full length PARP2. (A) wild type PARP2 and PARP2$^{S8,73A}$ mutant; (B) PARP2$^{S8A}$ mutant, PARP2$^{S73A}$ mutant; (C) PARP2$^{SA}$ mutant.

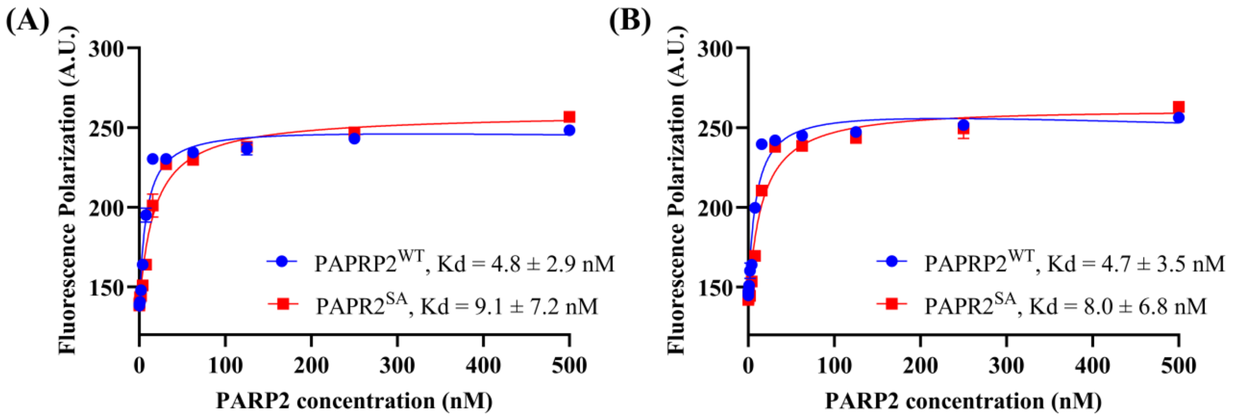


**Figure S2.** Fluorescence polarization-based Kd determination for oligo and PARP2 wild type (Blue) / no ser (Red). (A) Without HPF1 and (B) in the presence of 5 μM HPF1. All the reactions were performed in triplicates, and the data is shown mean ± s.d. of 3 independent experiments.

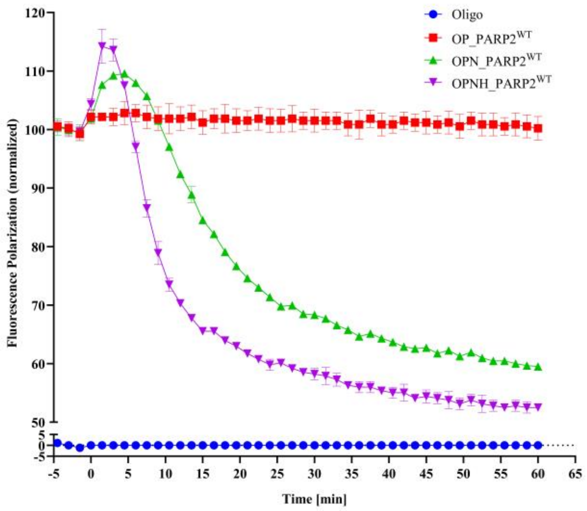


**Figure S3.** Representative fluorescence polarization assay to monitor the effect of HPF1 on release of wild type PARP2 from DNA. Data shown is average ± s. d. from three replicates and normalized with oligo alone (minimum FP) and DNA bounded PARP2 (maximum FP). In the presence of HPF1, PARP2 releases faster than without HPF1. [O: Oligo; OP: Oligo + PARP2; OPN: Oligo + PARP2 + $NAD^+$; OPNH: Oligo + PARP2 + $NAD^+$ + HPF1].

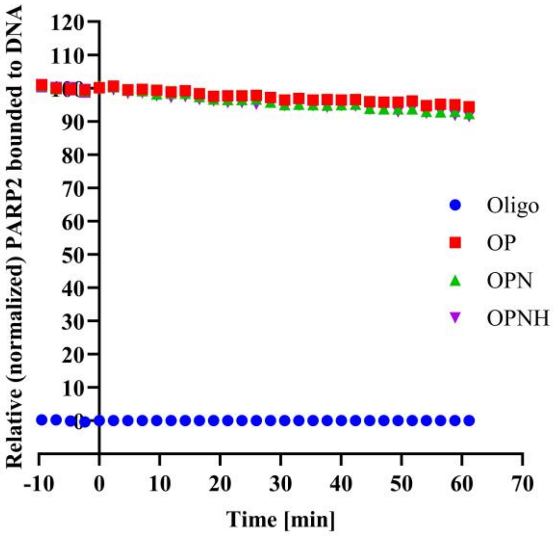


**Figure S4.** Representative fluorescence polarization assay to monitor the effect of PARP2 catalytic inhibitor on its release from DNA. Data shown is average ± s.d. from three replicates and normalized with oligo alone (minimum FP) and DNA bounded PARP2 (maximum FP). EB47 abolishes PARP2 catalytic activity, leading towards no PARylation of PARP2 which results in no change in FP signal. [O: Oligo; OP: Oligo + PARP2; OPN: Oligo + PARP2 + $NAD^+$; OPNH: Oligo + PARP2 + $NAD^+$ + HPF1].

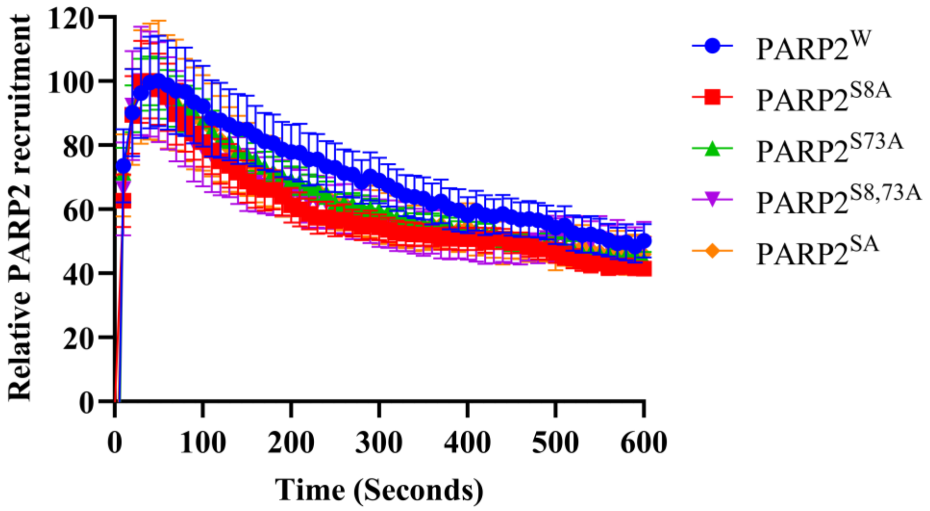


**Figure S5.** Relative kinetic recruitment and release of GFP-PARP2 from DNA lesion in PARP1/2 double knockout U2OS cells. Data shown as mean ± SEM (n > 5 per construct), normalized to pre-damage GFP intensity at micro irradiation site.

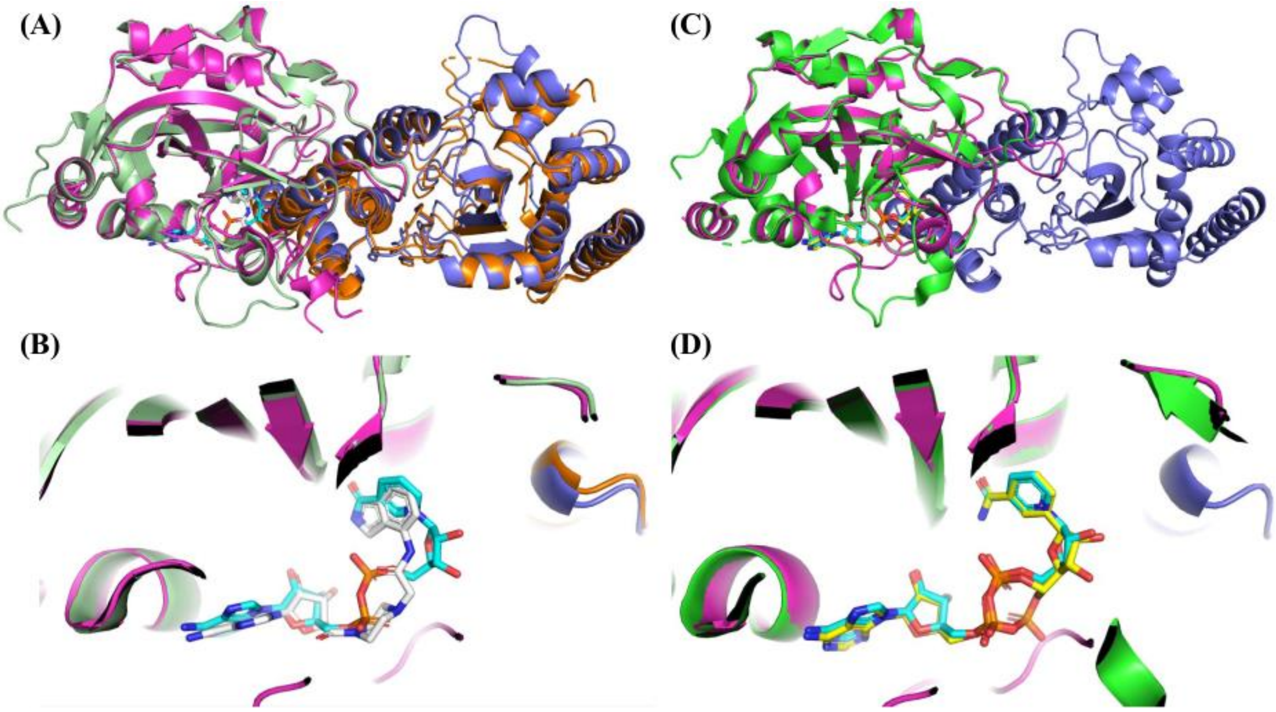


**Figure S6.** Structural analysis of Alphafold 3 generated PARP2-HPF1 complex and $NAD^+$ binding. (A) Structural alignment of crystal structure of PARP2 (ART)-HPF1 (PDB: 6TX3) with Alphafold generated model of PARP2 (ART)-HPF1, (B) binding analysis of modelled $NAD^+$ with crystalized inhibitor EB-47. (C) Structural alignment of PARP1 ART (PDB: 6BHV) with modeled PARP2-HPF1 complex, (D) binding analysis of modelled $NAD^+$ with crystalized PARP inhibitor and $NAD^+$ analog BAD. [PARP1 crystal structure (6BHV): PARP1 ART (green), BAD (yellow); PARP2-HPF1 crystal structure (6TX3): HPF1 (orange), PARP2 ART (pale green), EB-47 (grey); A3 model: HPF1 (blue), PARP2 ART (magenta), $NAD^+$ (cyan)].

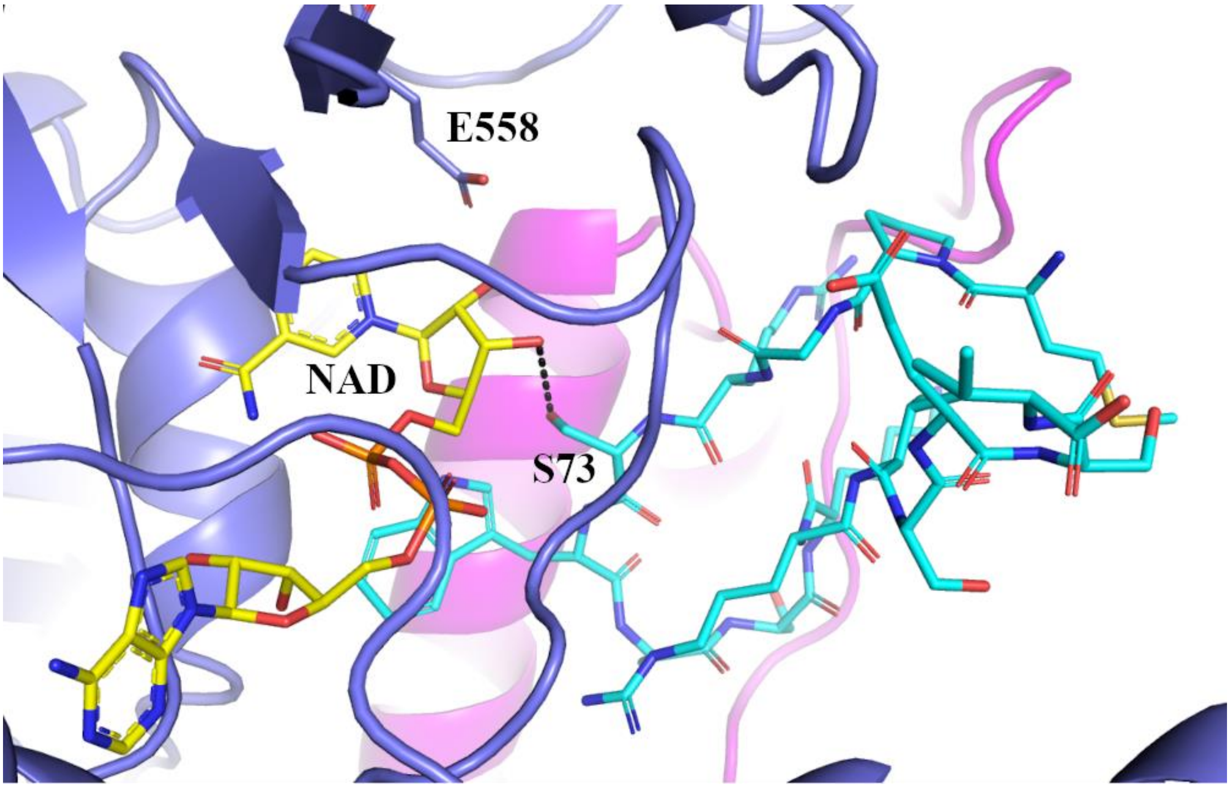


**Figure S7.** Alpha-fold 3 model of PARP2-HPF1-NAD$^+$ with N-terminal peptide complex. 15-mer peptide (cyan) interaction at PARP2 ART (blue)-HPF1 (magenta) where Ser73 is oriented near ribose sugar of NAD$^+$ (yellow). E558 is PARP2 (isoform-1) catalytic glutamate present near PARP(ART)-HPF1 junction.

**Table S1.** 15-mer peptide sequences from PARP2 isoform-1 which are used for modelling with PARP2-ART and HPF1 in Alphafold 3. Distances between "O" of serine side chain and O3' of $NAD^+$ ribose sugar are measured from the best scored model.

| Serine as target site | Amino acid number (start-end) | Peptide sequence | Distance (Å) between Ser side chain oxygen and $NAD^+$ O3' |
|---|---|---|---|
| S8 | 1-15 | MAARRRRSTGGGRAR | 3.4 |
| S20 | 13-27 | RARALNESKRVNNGN | 12.2 |
| S33 | 27-41 | NTAPEDSSPAKKTRR | 13.5 |
| S34 | 27-41 | NTAPEDSSPAKKTRR | 7.0 |
| S47 | 40-54 | RRCQRQESKKMPVAG | 6.6 |
| S73 | 69-83 | MPGRSWASKRVSESV | 3.9 |
| S76 | 69-83 | MPGRSWASKRVSESV | 12.7 |
| S80 | 75-89 | ASKRVSESVKALLLK | 10.6 |
| S82 | 75-89 | ASKRVSESVKALLLK | 12.6 |
